# Comparing the Statistical Fate of Paralogous and Orthologous Sequences

**DOI:** 10.1101/053843

**Authors:** Florian Massip, Michael Sheinman, Sophie Schbath, Peter F. Arndt

## Abstract

Since several decades, sequence alignment is a widely used tool in bioinformatics. For instance, finding homologous sequences with known function in large databases is used to get insight into the function of non-annotated genomic regions. Very efficient tools, like BLAST have been developed to identify and rank possible homologous sequences. To estimate the significance of the homology, the ranking of alignment scores takes a background model for random sequences into account. Using this model one can estimate the probability to find two exactly matching subsequences by chance in two unrelated sequences. The corresponding probability for two homologous sequences is much higher allowing to identify them. Here we focus on the distribution of lengths of exact sequence matches in protein coding regions pairs of evolutionary distant genomes. We show that this distribution exhibits a power-law tail with exponent α = —5. Developing a simple model of sequence evolution by substitutions and segmental duplications, we show analytically that paralogous and orthologous gene pairs contribute differently to this distribution. Our model explains the differences observed in the comparison of coding and non-coding parts of genomes, thus providing with a better understanding of statistical properties of genomic sequences and their evolution.

## I. INTRODUCTION

One of the first and most celebrated bioinformatic tools is sequence alignment [1, 17, 24]. Even today, the development of algorithms that are able to search for similar sequences of a query sequence in a huge database is still an active research field. In this matter, one needs to be able to distinguish sequence alignments that are due to a biological relatedness of two sequences from those which occur by random. Let us, for simplicity, disregard mismatching nucleotides and insertions and deletions (indels or gaps) in an alignment and only consider so-called maximal exact matches, i.e. sequences that are 100% identical and cannot be extended on both ends. In this case, the length distribution of matches is equivalent to the score distribution and can easily be calculated for an alignment of two random sequences where each nucleotide represents an i.i.d. random variable. We expect the number of matches to be distributed according to a geometric distribution, such that the number, *M*(*r*), of exact maximal matches of length *r* is given by

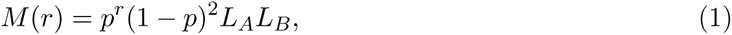

where *L*_*A*_ and *L*_*B*_ are the lengths of the two genomes, *p*^*r*^ is the probability that *r* nucleotides match, and (1—*p*)^2^ is the probability that a match is flanked by two mismatches. Here *p* = Σ_α_*ƒ*_*A*_(α)*ƒ*_*B*_(α), where *ƒ*_*x*_(α) is the frequency of nucleotide α in the genome of species *X* and the sum is taken over all nucleotides. Thus, the number of matches for a given length *r* is expected to decrease very fast as the length *r* increases, and for generic random genomes of hundreds of Mbp, one does not find any match longer than 25 bp.

For long matches, real genomes strongly violate the prediction of Eq. (1) due to the evolutionary relationships between and within genomes [21]. Comparing the genomes of recently diverged species, one finds regions in the two genomes that have not acquired even a single substitution. In the following, substitution refers to any genomic change which would disrupt a 100% identical match (as for instance a nucleotide exchange, an insertion, or a deletion). As the divergence time between the two species increases, such matches will get smaller very fast and most remaining long matches will be found either in exons or in ultraconserved elements [2], that both evolve under purifying selection.

Computing the match length distribution (MLD) from the comparison of human and mouse genomes, we thus expect to observe much longer exact matches than in a comparison of two random sequences of the same lengths. The observed MLD for exons and non-coding sequences in the human and mouse genomes is shown in Fig. 1. At the left end of the distribution, i.e. for *r* < 25 bp, the distribution is dominated by random matches, as described by Eq. (1). As expected, this MLD deviates from the random model for matches longer than 25 bp.

**Figure 1:**
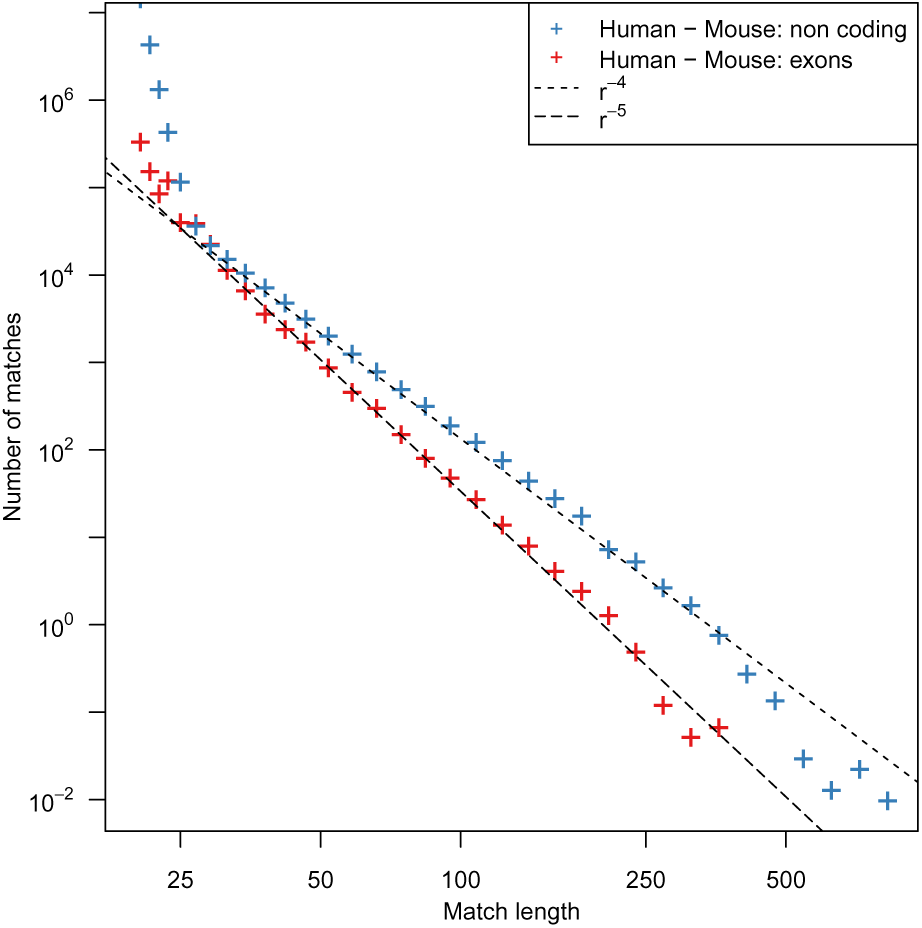
Two MLDs computed from the comparison of different subsets of the Human and the Mouse genomes. One MLD was computed from the comparison of the RepeatMasked non-coding part of both genomes (blue crosses) and the second is the result of the comparison of the coding part of both genomes (red crosses). Dashed lines represent power-law distributions with exponents α = — 4 and α = —5. All data are represented using a logarithmic binning to reduce the sampling noise [18], see the Materials and Methods section in the Supplementary Material for more details.

Interestingly, in this asymptotic regime, the MLD exhibits a power-law tail *M*(*r*) ~ *r*^α^ (identified as a straight line in the double logarithmic plot), where the exponent α is close to —5 for exonic sequences. This is in contrast to the MLD of non-coding sequences, where the exponent α is close to —4 [9, 10, 16]. This property appears to be impressively reproducible in the comparison of various pairs of species (see Fig S1). The value of the exponent α was calculated using the maximum likelihood estimator. To assess the robustness of this estimator, we also performed a bootstrap analysis that showed good agreement with the estimated value of α See Section V in the supplementary material. Discrepancies from the of power-law behavior can be observed for very long matches, since such matches are scarce and random noise distorts the distributions.

It is tempting to speculate that this peculiar behavior of exonic sequences is a direct consequence of their coding function and might reflect structural or other constrains of proteins. In the following, we demonstrate that this distribution can be explained by a simple evolutionary model that takes into account the generation of paralogous sequences (due to segmental duplications) and orthologous sequences (due to speciation) [6]. Further, we assume that paralogous and orthologous matches are subsequently broken down by random substitutions. This dynamics can be modelled by a well-known stick-breaking process [13, 26] introduced below. Since our model describes the existence of long matching sequence segments in two genomes, it also has to include selection. However we model selection in a minimal way, since we only assume that regional substitution rates are distributed, such that there are regions that evolve very slowly. Our model can therefore be viewed as a minimal model for evolution of functional sequences, which reproduces certain statistical features of their score distributions. We proceed now with the detailed introduction of the model.

## II. THEORY

### The Stick-Breaking Model

— Before we turn to the detailed description of our model, let us shortly introduce some relevant results on random stick-breaking. Consider a stick of length *K* at time *t* = 0, which will be sequentially broken at random positions into a collection of smaller sticks. Breaks occur with rate *μ* per unit length. The distribution of stick lengths at time *t*, denoted by *m*(*r,t*), follows the following integro-differential equation [15, 26]:

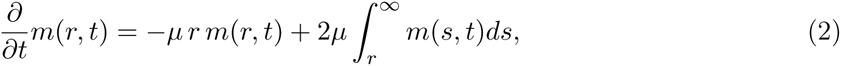

where the first term on the right hand side represents the loss of sticks of length *r* due to any break in the given stick and the second term represents the gain of sticks of length *r* from the disruption of longer sticks. Note that for any stick of length *s* > *r*, there are two possible positions at which a break would generate a stick of length *r*.

The initial state is one unbroken stick of length *K*, i.e. *m*(*r*, 0) = δ(*K,r*). The corresponding time-dependent solution is [26]:

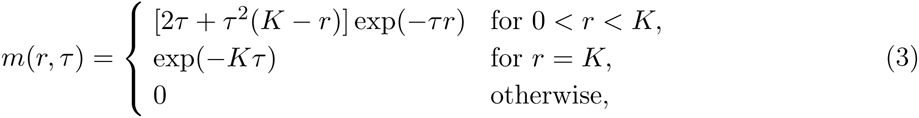

where we define the rescaled time *τ* = *μt*. Apart from the singularity at *r* = *K*, which accounts for the possibility that the stick is not even broken once, the distribution is dominated by an exponential function, i.e. there are far more small sticks than long ones. The average stick length is given by 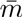(*τ*) = *K*/*τ*.

### The Match Length Distribution of Evolving Sequences

— The above stick-breaking process can be used to describe the break down of a long DNA match into several smaller ones by substitutions in either one of the two copies of the match. In a comparison of two species, A and B, long identical segments are the signature of homology relationships between the two sequences. These homologous sequences result either from the copy of the genetic material during the time of speciation, and are then orthologuous sequences (see the blue dashed line in Fig. 2) or due to segmental duplications in the ancestral genome, i.e. paralogous sequences (see the red dashed line in the same figure) [6].

**Figure 2:**
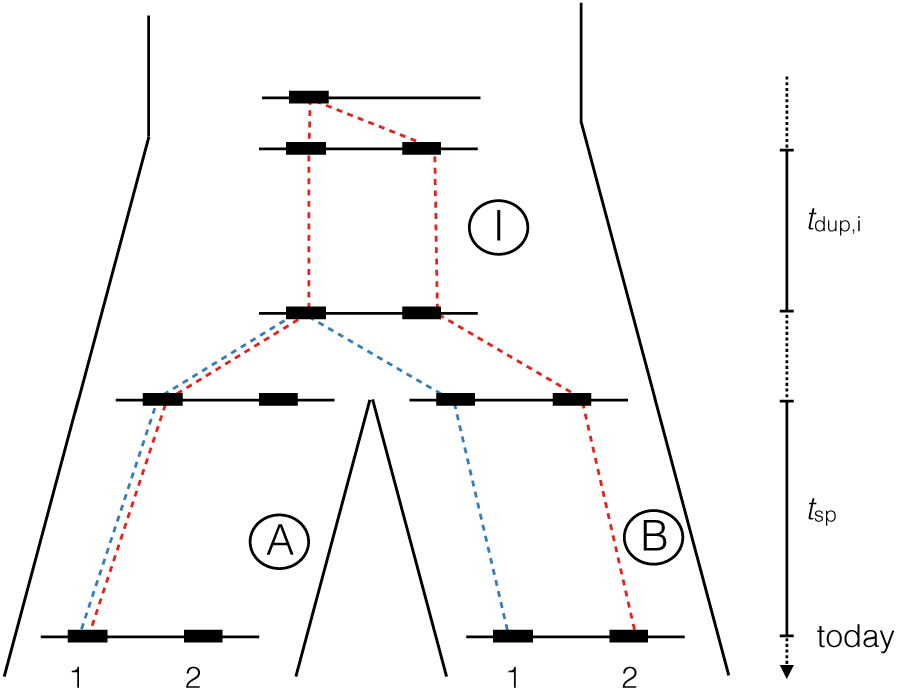
The different contributions to the match length distribution. Sequence 1 was duplicated in the ancestral species I. Fhis duplication gives rise to two paralogous sequence pairs: sequence 1 in A with sequence B 2 in B (red dashed line), and sequence B 1 and sequence A 2. Sequence 1 in A is orthologous to sequence 1 in B (blue dashed line), and Sequence 2 in A is orthologous to sequence 2 in B. For clarity, we highlight only one pair only for each exemple on the figure.

The match length distribution (MLD) is then given by the integral

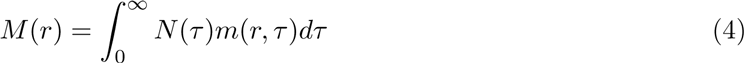

where *N*(*τ*) is the number of homologous sequences with divergence *τ* and *m*(*r,τ*) is given in Eq. (3), see also Ref.[16]. The divergence between a pair of orthologous sequences is the sum of two contributions *τ* = *μ*_*A,i*_ *t*_sp_ + *μ*_*B,i*_ *t*_sp_, where *t*_sp_ is the time since the two species diverged and *i* is an index for regions in the genomes. The regional mutation rates *μ*_*A,i*_ and *μ*_*B,i*_ in the two species are themselves distributed and assumed to be independent from each other. We therefore define *N*_*AB*_

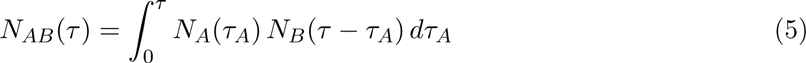

where *N*_*A*_(*τ*) (resp. *N*_*B*_(*τ*)) is the number of sequences with divergence *τ* from the last common ancestor *I* in species *A* (resp. *B*). However if the two regions are paralogous, the divergence *τ* is a sum of three independent contributions *τ* = *μ*_*A,i*_*t*_sp_ + (*μ*_*I,i*_ + *μI,j*)*t*_dup_ + *μ*_*B,j*_ *t*_sp>_where *t*_dup_ represent the time elapsed between the segmental duplication and the split of the two species. There are *N*_*AIB*_(*τ*) paralogous sequences with divergence *τ*, with

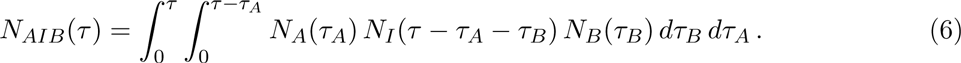

For our purposes we are not interested in the full functional form of the distributions in (Eqs. (5) and ((6) but only have to consider their behavior for small *τ* →0, because long matches (and thus the tail of the distribution of the match length distribution *M*(*r*)) stem from homologous exons that exhibit a small divergence *τ*. A more general discussion about the functionnal form of the distrbution of pairwise distances can be found in ref.[23]. We therefore take the Taylor expansion of the distributions *N*(*τ*) around *τ* = 0. Using Leibniz’s formula to take the derivative under the integral sign [7] we find for orthologous exons (see details in the Supplementary Material)

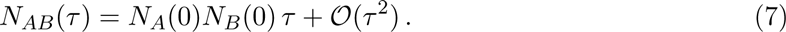

and subsequently the match length distribution [16]

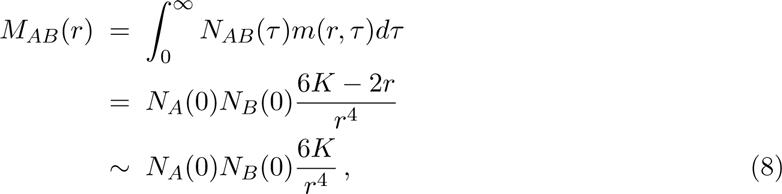

as *K* is much larger than *r*. In contrast, expanding (Eq. (6) around *τ* = 0, one can see that for paralogous pairs, the number of regions with divergence *τ* increases as *τ*^2^ in the small *τ* limit (see details in the Supplementary Material)

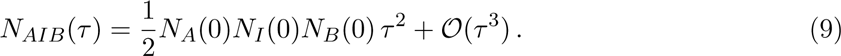

Therefore the match length distribution exhibits a power-law tail with exponent α = —5:

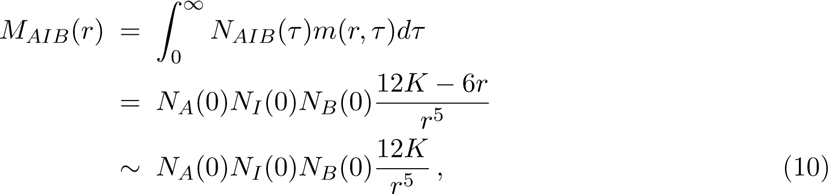

as *K* is much larger than *r*.

Depending on the number of orthologous sequences Q_orthoiog_ and paralogous sequences Q_paraiog_, we will be able to distinguish two regimes: one where the MLD follows an α = —4 power-law and one where it follows an α = —5 power-law. From Eqs. (8) and (10), it is straightforward to find that the cross-over point *r*_*c*_ between those regimes (see Fig. 3) is at

**Figure 3:**
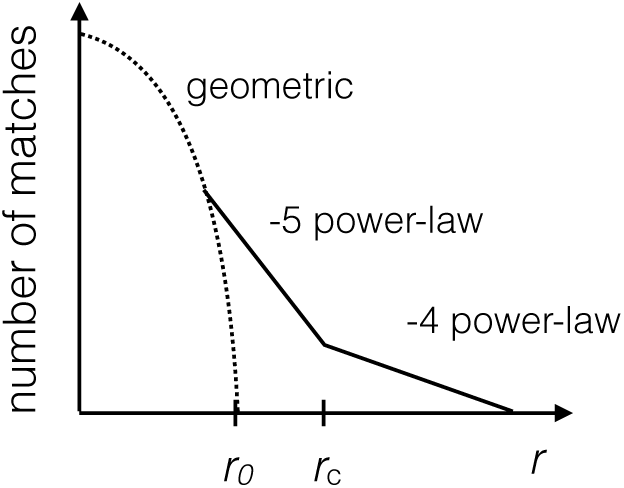
Schematic drawing of the match length distribution in a double logarithmic plot. The two regimes exhibiting a —4 and —5 power-law (continuous lines) are separated by a cross-over point. For very small match lengths the geometric distribution due to random matches, see Eq. (1), dominates (dotted line).

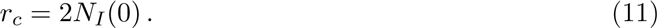

Recall that *N*_*I*_(0) is defined as the number of paralogous segments, at the time of the split, that have not mutated even a single time since the duplication event. Thus, this term is proportional to the ratio of the duplication rate over the mutation rate. If *N*_*I*_(0) ≫ 10, there are significantly more paralogous sequences compared to orthologous ones and the cross-over value, *r*_*c*_, is large. Then, only the α = —5 power-law tail will be observed. On the other hand, if *N*_*I*_(0) ≪10, then the crossing point *r*_*c*_ is expected to be smaller than 20 such that the α = —5 power-law only holds for lengths were the distribution is anyway dominated by random matches. In contrast to previous models[16], this model does take into account the contribution of paralogous sequences, and can explain both power-law behaviors and therefore predicts the crossing point between the two regimes.

## III. NUMERICAL VALIDATION

Our theoretical considerations predict a complex behavior of the match length distribution under the described evolutionary dynamics. The key ingredients are segmental duplications, generating paralogous sequences in an ancestral genome, and point mutations, that break identical pairs of paralogous and orthologous sequences of the two genomes after speciation into smaller pieces. To illustrate our theoretical predictions concerning the two power-laws, as well as the existence of the cross-over point *r*_*c*_, we simulated the evolution of sequences according to the discussed scenario.

We describe the evolution of a genome of length *L* according to two simple processes, point mutations and segmental duplications. Point mutations exchange one base pair by another one and occur with rate *μ* per bp and unit of time. Note that to mimic the existence of regions under different degrees of selective pressure, we allow for regional differences of the point mutation rates. Segmental duplications copy a contiguous segment of *K* nucleotides to a new position where it replaces the same amount of nucleotides, such that the total length of the genome stays constant. Segmental duplications occur with rate λ per bp.

The evolutionary scenario of our simulation has two stages (see Fig. 2). At time *t* = 0, we generate a random iid sequence *S*. During a time *t*_0_, this sequence evolves according to the two described processes. In this first stage, the mutation rate is the same for all positions. At the end of this stage, the sequence represents the common ancestral genome of two species. At the beginning of the second stage, we copy the entire sequence of the common ancestor to generate the genomes of the two species *A* and *B*. These sequences are then subdivided into *M* continuous regions of equal length. In each such region *j*, the point mutation rates *μ*_*Aj*_ (resp. *μ*_*B,j*_) are the same for all sites *i* and are drawn from an exponential distribution with mean *μ* (i.e. the point mutation rate during the first stage). The exponential distribution stipulates the least information under the given constraints. For more details about the simulation procedure, see Section V in the Supplementary Material.

We show the result of of the comparison of simulated sequences on the left panel of Fig. 4. One can clearly identify a power-law tail in the match length distribution, which for match length 20 < *r* < 100 has an exponent α= —5 and an exponent α= —4 for longer matches *r* > 100. For simulated sequences, we can also easily classify homologous sequences into orthologous and paralogous sequences (while for natural sequences, paralogs and orthologs are not as easy to distinguish due to genomic rearrangements). Thus, we can also compute the MLD obtained from the comparison of paralogs, and the MLD obtained from the comparison of orthologs for simulated sequences. We show these two different distributions on the right panel of Fig. 4, where one can clearly observe that orthologous sequences generate an α= —4 power-law distribution while paralogous matches generate an α = —5 power-law distribution. In such a plot, one can further easily identify the crossing point as the value of *r* for which we obtain more matches from the comparison of orthologs than from the comparison of paralogs.

**Figure 4:**
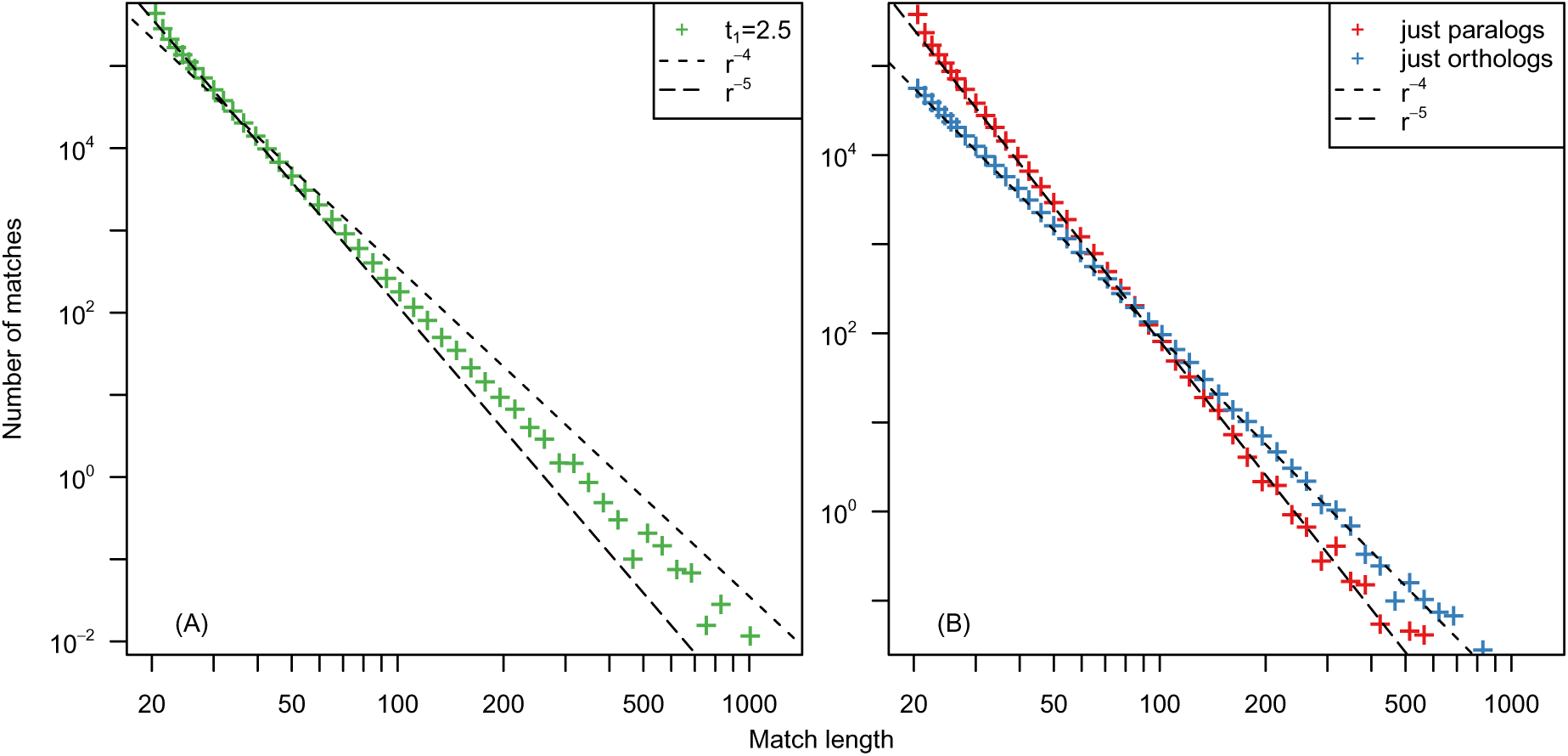
The MLD computed from 1000 simulated sequences according to the procedure described in the main text. Data are represented using the logarithmic binning. On the left panel, we show the MLD generated computed from all possible matches, while on the right panel, we represent two different MLDs: one computed from paralogous matches only (red), and one computed from the orthologous matches only (blue). One can see that the two different MLDs crosses close to the expected crossing value *r*_*c*_ = 100.

In the previous section the value of this crossing point between the two regimes was predicted to be *r*_c_ = 2*N*_*I*_(0) see (Eq. (11), where *N*_*I*_(0) is the number of paralogous segments with a divergence *τ* = 0 just before the species split. In our simulation procedure, *N*_*I*_(0) is simply the number of sequences that have been duplicated but that have not been mutated yet at the splitting time *t* = *t*_0_. This number is known to be *N*_*I*_(0) = 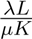[15]. In our simulations, we have chosen λ = 0.05, *μ* = 1, *K* = 1000 and *L* = 10^7^ and therefore the crossing over is predicted to happen at *r*_*c*_ = 100, which is in good agreement with our observations in Fig. 4. Thus, the results of our simulations are in good agreement with our analytical predictions.

### IV. DISCUSSION

We developed a simple model that accounts for power-law tails in the length distribution of exact matches between two genomes. Our model assumes regional differences of the selective pressure such that the substitution rates in a region are drawn from a certain distribution. However for naturally evolving exons the selective pressure varies also on shorter length scales. For instance, some nucleotides for many codons can be synonymously substituted by another one, mostly at third codon positions. Therefore, these substitutions at 3rd codon positions occur with a higher rate than non-synonymous ones. Hence, exons are expected to break preferentially at positions 3*n*, with *n* ∊ ℕ, such that the matches with 100% identity would have lengths 3*n* + 2 with integer *n*. Classifying genomic matches according to the remainder which is left when dividing their length by 3, we observe an almost 10-fold over-representation of matches with length 3*n* + 2 over matches of lengths 3*n* and 3*n* + 1, see Fig. 5. This suggests that the match-breaking process is dominated by the synonymous mutation rate which does not seem to vary significantly along an exon, while it does vary from one exon to another.

**Figure 5:**
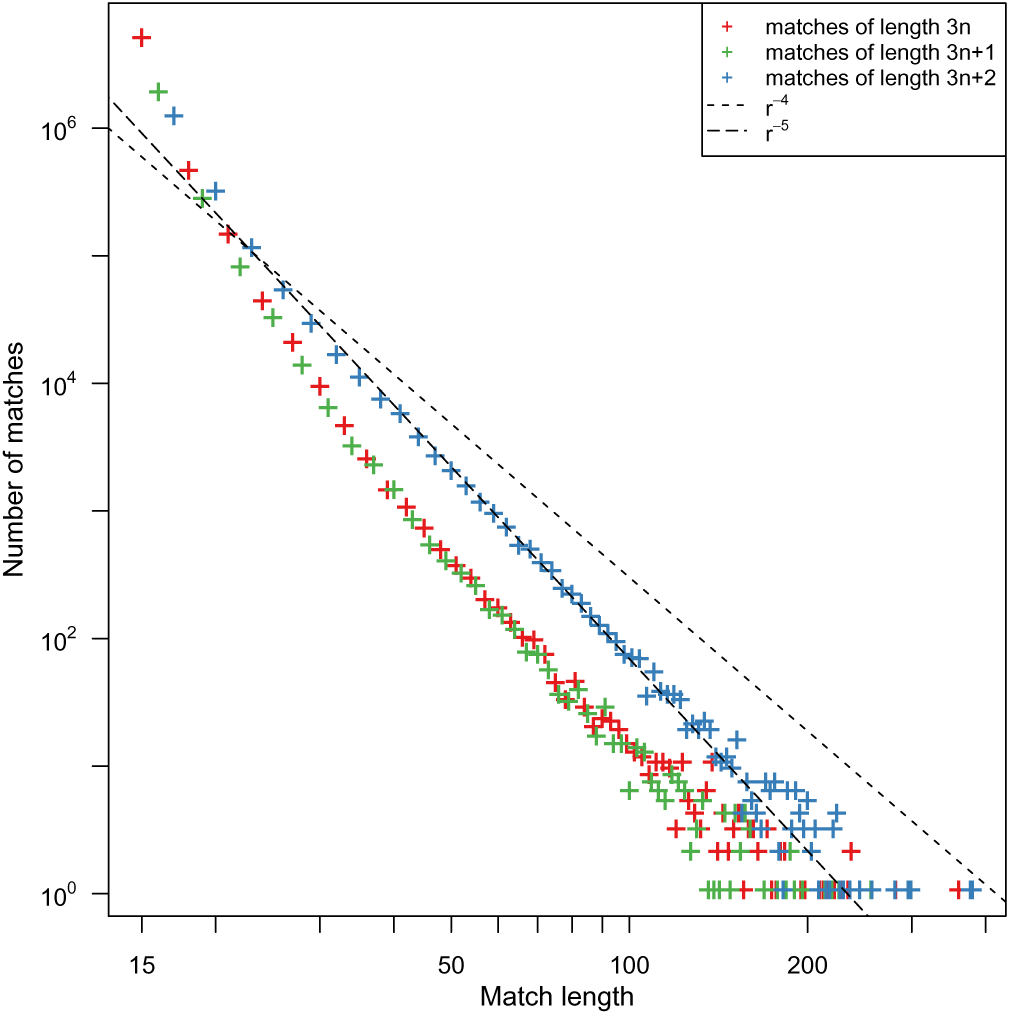
The MLD computed from the comparison of human and mouse exome, represented without logarithmic binning. 3 different colors are used to represent matches of length 3*n*, 3*n*+1 and 3*n*+2. Dashed lines represent power-law distribution with exponent α = −4 and α = −5.

Using the presented model, the puzzling observation of an α= —4 power-law tail in the MLD in the comparison of the human and mouse genomes and a corresponding α= —5 power-law tail in the comparison of their exomes can be explained. Although the sequences stem from the same species, the relative amount of paralogous to orthologous sequence segments is different in the two data sets, which subsequently leads to different cross-over point *r*_*c*_. Because of the selective pressure on coding exons, the number of non-mutated paralogous sequences at the time of species divergence *N*_*I*_(0) is higher (relative to the number of orthologous sequences) in the exonic dataset than in the non-coding dataset. Thus, the cross-over point in exomes *r*_*c*_ is larger than the longest observed match and only the α= —5 can be observed.

The opposite is true for matches in the alignment of non-coding sequences. Quantitatively, in this set, paralogous sequences play a lesser role and therefore only the α= —4 power-law is observed (see Fig. 1). This is surprising, as the duplication rate is thought to be roughly the same in the coding and non-coding parts of genomes. To confirm this paradoxical observation, we classified matches according to the uniqueness of their sequences in both genomes. Assuming that unique matches are more likely to be orthologous, this gives us a rough classification of homologs into orthologs and paralogs, although matches unique in both sequences can be paralogs. After the classification of all matches, our analysis made apparent that matches unique in both genomes dominate the MLD in the comparison of the non-coding parts of the genomes, while matches with several occurrences in either (or both) genomes dominate the distribution in the case of the comparison of exomes (see Fig. S2 in the Supplementary Material). Moreover, we computed the MLD from the set of non-unique matches of the non-coding part of genomes. In this comparison, the contribution of paralogs is expected to be much higher than in the full set. As expected, this MLD also exhibits an α = —5 power-law (see Fig. S2) in the Supplementary Material), confirming that the relative contribution of orthologs and paralogs is responsible for the shape of the MLD. These differences in the proportion of paralogous sequences in the coding and non-coding DNA is likely due to the fact that paralogs are more often retained in coding sequences than in the non-coding part of genomes. Since there are much more non-coding sequences in both genomes, we also observe at least 10 times more matches in the comparison of non-coding sequences than in the comparison of exomes.

The presented model does not account for changes in the divergence rates after a duplication, a phenomenon which is well documented following a gene duplication[11, 19, 20, 22]. To assess the impact of this phenomenon on the MLD, we performed simulations where the two paralogous segments are assigned different and independent mutation rates. Interestingly, these simulations yield qualitatively similar results than the simpler model introduced above (see Fig. S3). This new condition does not affect the value of the number of paralogous sequences that have not diverged at the time of the split (i.e. the value of *N*_*I*_(0)), and thus the shape of the distribution.

The model we present is very simple, and more realistic models of genome evolution include many more evolutionary processes [5]. For instance one could include a transition/transversion bias in the mutational process, variations of mutation rates in time, a codon usage bias or different rates of duplications within and between chromosomes. However, since in the end we just consider identical matching sequences (and do not differentiate between miss-matches due to transitions and transversions) and want to explain the power-law tail in the MLD, all these additional model details are not expected to affect the results.

In this paper, we demonstrate that on the genome-wide scale, the length distribution of identical homologous sequence segments in a comparative alignment is non-trivial and exhibits a power-law tail, and we propose a simple model able to explain such distributions. While paralogous sequences, which had been duplicated before the species diverged generate a power-law tail with exponent α= —5, orthologous sequences generate a power-law tail with exponent α= —4. Depending on the relative amount of paralogous to orthologous sequences there is a cross-over between these two power-law regimes. The exponent of the power-law tail in the comparative MLD can therefore be a litmus test for the abundance of paralogous relative to orthologous sequences, while it is usually difficult to distinguish between orthologous and paralogous sequences using classical bioinformatic methods [4, 8, 25]. If paralogous sequences dominate (as for instance when comparing the exomes of human and mouse) the cross-over happens for long matches and the apparent exponent is equal to −5, otherwise it is equal to −4 (as for instance when comparing the non-coding part of human and mouse genomes).

Our method is very easy to apply. In particular, it does not require that genomes are fully assembled as long as the continuous sequences are longer than 1kbp, comparable to the longest matches one expects. A natural extension of our method would be to apply it to sequences from metagenomic samples in order to assess relative amounts of paralogous and orthologous sequences. However, in this setting also horizontal gene transfer, which is common among prokaryotes and generates homologous sequence segments even between unrelated genomes, has to be included into the model. Our computational model can be easily be extended to take into account these and other more complex biological processes, using for instance already developed tools [5], which would allow to assess their impact on our results. The analysis of these models will be the subject of future work.

This study shows that even very simple models can often successfully be applied to seemingly very complex phenomena in biology. We were able to present a minimal model for the evolution of homologous sequences that includes effects due to segmental duplications and evolution under selective constrains — the two processes that are responsible for a power-law tail in the length distribution of identical matching sequences.

## V. MATERIAL AND METHODS

### Genomes

— Unless otherwise stated, all genomes and their specific annotations (such as repeat elements and exons) were downloaded from the `ensembl` website [3] using the Perl API (version 80); the corresponding release of the Human genome is GRCh37. In all cases, we downloaded the RepeatMasked versions of genomes available in `ensembl` databases.

### Computing MLDs

— To find all identical matches (both in the case of self and comparative alignments), we used the `mummer` pipeline [14] (version 3.0), which allows to find all maximal exact matches between a “query” and a reference sequence using a computationally efficient suffix tree approach. For our analyses, we used the following options:

- `-maxmatch` such that `mummer` searches for all matches regardless of their uniqueness (by default, only matches unique in the query sequence are retrieved).
- `-n` that states that only ‘A’, ‘T’, ‘C and ‘G’ can match (any other character always results in a mismatch).
- `-b` such that `mummer` searches for matches on both strands. To do so, the reverse complement sequence of the query file is computed, and `mummer` searches matches between the two forward sequences, as well as matches between the forward sequence of the reference and the reverse complement sequence of the query.
- `-l 20` to filter out matches smaller than 20 bp.

`mummer`’s output consists in a file with three columns representing for each match its position in the query sequence, its position in the reference sequence and its length. The number of matches expected for a random iid sequence grows quadratically with *L*. For instance one expects 3.5 10^18^ matches of length 2 in a comparison of two sequences of length *L* = 10^9^ bp (see Eq. (1)). This explains why one has to define a threshold for the length of matches that `mummer` should output especially when comparing entire eukaryotic genomes. Depending on the length of the sequences to compare, we might vary the value of this threshold.

### Logarithmic Binning

— Power-laws appear in the tail of distributions, meaning that they are associated to rare events, which are thus subject to strong fluctuations. The high impact of noise in the tail of the distribution can make the assessment of the exponent of the distribution difficult. A way to resolve this issue is to increase the size of the bins with the value of the horizontal axes and renormalise the data accordingly. Namely, the observed values for each bin are divided by the size of the bin. The most common choice to do this is known as the logarithmic binning procedure, which consists in increasing the size of the bin by a constant factor. Note that by doing so, one dramatically reduces the number of data points and some information is lost as one aggregates different data points together in the same bin (and data with different values are summarised together as one data point). Therefore it is often useful to consider both representations, with and without the logarithmic binning; for instance to observe the overrepresentation of matches with length 3*n* + 2 for integer *n* no logarithmic binning should be used. See reference [18] for more details on the logarithmic binning procedure, and on power-law distributions.

### Simulating the Evolution of DNA Sequences

— To simulate our evolutionary models, we proceeded as follows. A sequence of nucleotides *S* = (*s*_1_,…,*s*_*L*_) of length *L* with *s*_*i*_ ∊ {A, C, G, T} is evolved through time in small time intervals Δt. The time intervals Δt are small enough such that for all considered evolutionary processes *E* of our model, which are assumed to occur with rate *ρE*, we have *ρELΔt* ≪ 1. At each step, random numbers 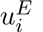 for all positions *i* and possible evolutionary processes *E* are drawn from a uniform distribution. The event *E* then occurs at position *i* if 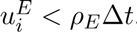. These steps are repeated until the desired time *t* has elapsed.

Sequences evolve according to two simple processes, point mutations and segmental duplications. Point mutations exchange one nucleotide by another and occur with rate *μ* per bp and unit of time. Note that to mimic the existence of regions under different degrees of selective pressure we allow for regional differences of the point mutation rates. Segmental duplications copy a contiguous segment of *K* nucleotides starting at position *c* and paste them to a different position *v*, such that the *K* nucleotides at positions *v* to *v* + *K* - 1 are replaced by the ones from position *c* to *c* + *K* - 1. As a consequence, the total length of the sequence *L* stays constant in time. The segmental duplication process occurs with rate λ per bp and per unit of time.

The evolutionary scenario of our simulation has two stages, as shown in Fig. 2. At time *t* = 0, we start with a random iid sequence *S* with equal proportions of all 4 nucleotides. During a time interval of length *t*_*0*_, this sequence evolves according to the two described processes. In this first stage, the mutation rate is the same for all positions. At the end of this stage, the sequence represents the common ancestor of the two species.

At the beginning of the second stage, we duplicate the entire sequence of the common ancestor to generate the genomes of the two species *A* and *B*. These sequences are then subdivided into *M* continuous regions of equal length, in which the point mutation rates *μ*_*A,j*_ (resp. *μ*_*B,j*_) are the same for all sites in a region *j* and are drawn from the same exponential distribution of mean *μ*, i.e. the point mutation rate during the first stage. The exponential distribution stipulates the least information under the given constraints.

For simplicity, the length of the *M* continuous regions is set to *K* and the segmental duplication rates in both species λ during the second stage are set to zero. Both species then evolve independently for a divergence time *t*_sp_, and we compute the MLD from a comparison of the sequences of the two species *A* and *B*. Note that even when we chose finite duplication rates after the split (i.e. λ > 0 in the second stage), the MLDs obtained from the simulated sequences were not qualitatively different.

To control for the potential impact of our choice to keep the genome size constant on our results, we also simulated the evolution of sequences where duplicated segments where added to the sequences (thus generating growing genomes). In that case, duplicates were added at the very end of the sequence, such that duplicates do not disrupt pre-existing matches. This control experiment yields qualitative similar results, in agreement with our theoretical considerations (data not shown).

### Simulating the Evolution of DNA Sequences

— To simulate our evolutionary models, we proceeded as follows. A sequence of nucleotides *S* = (*s*1,…, *s*_*L*_) of length *L* with *s*_*i*_ ∊ {A, C, G, T} is evolved through time in small time intervals Δt. The time intervals Δt are small enough such that for all considered evolutionary processes *E* of our model, which are assumed to occur with rate *ρE*, we have *ρELΔt* ≪ 1. At each step, random numbers 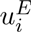 for all positions *i* and possible evolutionary processes *E* are drawn from a uniform distribution. The event *E* then occurs at position *i* if 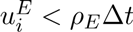. These steps are repeated until the desired time *t* has elapsed.

Sequences evolve according to two simple processes, point mutations and segmental duplications. Point mutations exchange one nucleotide by another and occur with rate *μ* per bp and unit of time. Note that to mimic the existence of regions under different degrees of selective pressure we allow for regional differences of the point mutation rates. Segmental duplications copy a contiguous segment of *K* nucleotides starting at position *c* and paste them to a different position *v*, such that the *K* nucleotides at positions *v* to *v* + *K* — 1 are replaced by the ones from position *c* to *c* + *K* — 1. As a consequence, the total length of the sequence *L* stays constant in time. The segmental duplication process occurs with rate λ per bp and per unit of time.

The evolutionary scenario of our simulation has two stages, as shown in Fig. 2. At time *t* = 0, we start with a random iid sequence *S* with equal proportions of all 4 nucleotides. During a time interval of length *t*_0_, this sequence evolves according to the two described processes. In this first stage, the mutation rate is the same for all positions. At the end of this stage, the sequence represents the common ancestor of the two species.

At the beginning of the second stage, we duplicate the entire sequence of the common ancestor to generate the genomes of the two species *A* and *B*. These sequences are then subdivided into *M* continuous regions of equal length, in which the point mutation rates *μ*_*A,j*_ (resp. *μ*_*B,j*_) are the same for all sites in a region *j* and are drawn from the same exponential distribution of mean *μ*, i.e. the point mutation rate during the first stage. The exponential distribution stipulates the least information under the given constraints.

For simplicity, the length of the *M* continuous regions is set to *K* and the segmental duplication rates in both species λ during the second stage are set to zero. Both species then evolve independently for a divergence time *t*_sp_, and we compute the MLD from a comparison of the sequences of the two species *A* and *B*. Note that even when we chose finite duplication rates after the split (i.e. λ > 0 in the second stage), the MLDs obtained from the simulated sequences were not qualitatively different.

To control for the potential impact of our choice to keep the genome size constant on our results, we also simulated the evolution of sequences where duplicated segments where added to the sequences (thus generating growing genomes). In that case, duplicates were added at the very end of the sequence, such that duplicates do not disrupt pre-existing matches. This control experiment yields qualitative similar results, in agreement with our theoretical considerations (data not shown).

### Estimating the value of the Power-law Exponent

To estimate the value of the exponent of the power-law α we compute the maximum likelihood estimator. The estimator 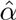 is simply the value of α that maximizes the log likelihood ***L***

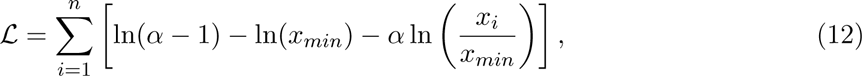

such that

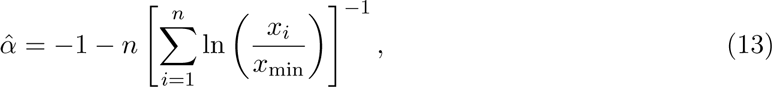

while the value of *x*_min_ has to be determined visually. This estimator is also sometimes referred to as the Hill estimator [12].

To estimate the robustness of the value of the exponnents found using this method, we proceeded to block boostrap experiments on the Human Mouse exome comparison. For each boostrap, we sampled 5% of the mouse exons, and compared them to all human exons. In each experiment, we calculated the exponnent of the MLD using the maximum likelihood estimator as described in [18]. We repeated this procedure a 100 times. Values of *α* were all in the range [—4.7, —5.2] and the mean value for the exponent was α = —4.9.

